# Solar Bird Banding: Notes on Changes in Avian Behavior While Mist-netting During an Eclipse

**DOI:** 10.1101/2023.12.09.570954

**Authors:** Adara DeNiro, Kyle D. Kittelberger, Atoosa M. Samani, Çağan Hakkı Şekercioğlu

## Abstract

Solar eclipses present rare celestial events that can elicit unique behavioral responses in animals, yet comprehensive studies on these phenomena, particularly concerning bird behavior, remain limited. This study, conducted at the Red Butte Canyon Research Natural Area in Utah during the annular solar eclipse on October 14, 2023, aimed to document and analyze avian activity using bird banding data. Leveraging 11 years of banding records, we observed a surprising positive peak in bird captures, indicating increased activity during the eclipse, challenging conventional expectations of decreased activity during peak totality. The unexpected, record-breaking captures on the eclipse day at this location, which also surpassed the average trend in captures over time for 18 other banding days in mid-October, highlights the complexity of bird behavior during celestial events. This study marks the first known published effort to conduct bird banding during a solar eclipse. Quantitative analyses, including species composition and capture trends, contribute to a nuanced understanding of avian responses to the eclipse. This study underscores the importance of empirical research in unraveling the intricacies of how birds navigate and adapt to unique environmental conditions created by solar eclipses.

## 1. INTRODUCTION

A solar eclipse is an uncommon phenomenon that occurs when the moon obscures the sun’s disk (Wiantoro 2019; Iskandar and Iskandar 2020), casting a shadow on Earth that produces a narrow path of totality (Nilsson et al. 2018; Brinley Buckley et al. 2018). Naturally, there is notable curiosity about the impact of solar eclipses on animal behavior (Wheeler et al. 1935; Pandey and Shukla 1982; Gil-Burmann and Beltrami 2003; Nilsson et al. 2018; Iskandar and Iskandar 2020). Due to the rare nature of this astronomical event (Stephenson 1982; Gil-Burmann and Beltrami 2003; Wiantoro 2019), this phenomenon is novel to those animals that experience its effects and, consequently, may induce anomalous behaviors. In many instances, animals appear to respond to solar eclipses in a manner consistent with their typical reactions to the onset of dusk (Wheeler et al. 1935; Wiantoro 2019). Diurnal species often exhibit a decline in activity, mirroring their natural response to approaching nighttime (Mousley 1933; Wheeler et al. 1935; Wiantoro 2019). Conversely, crepuscular or nocturnal species have shown an increase in activity during these celestial events (Dubrovsky and Tytar 2015; Brinley Buckley et al. 2018).

Birds can serve as a key model to study the effect of photic conditions on behavioral and physiological changes (Temple et al. 1989), especially since they are the best-studied group of organisms (Ali Tabur and Ayvaz 2010; Kittelberger et al. 2021) and there are continuous observations and research on birds (Kittelberger et al. 2023). For instance, several studies have noted unusual behavior amongst avian species in response to solar eclipses. During a solar eclipse in Central Ethiopia in 2020, there was a profound decrease in calling, singing, foraging and moving, and courtship behavior amongst the birds observed (Mekonen 2021). Likewise, some birds significantly altered their vocalizations during the 2017 eclipse in the United States (Brinley Buckley et al. 2018). In addition, on the north coast of Venezuela in 1998, birds ceased to forage over the water, and diurnal birds such as pelicans sought out their evening roosts during totality (Tramer 2000). Other observations parallel these sightings, such as in Central Sulawesi, Indonesia, on March 9^th^, 2016, where bird colonies exhibited a decrease in foraging during the initial stages of the solar eclipse, followed by swift disappearance during the period of totality (Wiantoro 2019). However, overall studies on the behavioral responses of animals, including birds, to eclipses are limited due to the infrequent and quick nature of eclipses (Nilsson et al. 2018), and what has been documented often is based on anecdotal observations rather than scientific ones (Brinley Buckley et al. 2018).

The annular solar eclipse took place on October 14, 2023, with the path of totality spanning North America, particularly across the western United States (Hurst et al. 2023). In Salt Lake City, Utah, the obscuration of the sun reached 86.78% (Time and Date 2023). Over the period from 2001 to 2023, North America experienced a total of 10 annular and total solar eclipses (Espenak 1987). Due to the infrequency of these celestial events, there are limited opportunities to comprehend the distinctiveness of biotic effects from each occurrence, given the variations in timing, locations, and atmospheric conditions. Consequently, this infrequency of solar eclipses in North America is compounded by a dearth of structured data collection, such as from bird banding or soundscape recordings, of birds in the field during these events to aid in our understanding of bird behavior during eclipses to changes in light conditions (but see Mendoza 2018; Nilsson et al. 2018; Brinley Buckley et al. 2018; Mekonen 2021).

In this study, our aim was to measure bird activity throughout an eclipse. Utilizing bird banding data gathered during the 2023 annular solar eclipse in Salt Lake City, Utah, we conducted a comparative analysis with data from days in the month of October at the same location across multiple years. This approach allowed us to examine avian behavior and movement patterns in response to the eclipse. We hypothesized that the eclipse would noticeably alter bird activity. Specifically, we predicted that bird captures would decline during peak totality and mirror bird behavior during the evening at dusk. This study provides the first known documentation of results from bird banding during an eclipse.

## 2. METHODS

### 2.1 Study Site

This research was conducted at the Red Butte Canyon Research Natural Area (RBC), a protected canyon located in the Wasatch Range of Utah, to the east of Salt Lake City and the University of Utah (Ehleringer et al. 1992; Barnick et al. 2022). This access-restricted site is only open to permitted researchers and has been protected since 1862 (Ehleringer et al. 1992; Barnick et al. 2022). Our bird banding station is located at Parleys Fork within the RBC around a central wetland at roughly 1736 m asl (40.787573, -111.796584), consisting of a wet meadow/marsh and several streams (including Red Butte Creek) surrounded by open montane forest and other riparian vegetation. Red Butte is located within the Western or Pacific flyway (La Sorte et al. 2014; Rosenberg et al. 2019).

### 2.2 Bird Banding

Bird banding has occurred annually at Red Butte from 2012 through 2023. Most annual banding seasons tend to last from late April through early November, encapsulating the entirety of spring and fall long-distance land bird migration in northeastern Utah, as well as the temperate avian breeding season. We used on-average thirteen 38 mm mesh mist-nets, each measuring 12 m by 3 m, with nets either been distanced from one another at each location or connected lengthwise to create a net wall. Typically, the nets were opened 30 minutes before sunrise and checked every half-hour for the subsequent six hours before closing, with the start and end times recorded. We also recorded the temperature at the opening and closing of each visit. Adjustments to the timing were occasionally made due to adverse weather conditions or other complications.

We extracted all birds from nets and first identified them to species. We then fitted aluminum butt-end leg bands (Bird Banding Laboratory, USGS) on the legs of newly-captured birds, and recorded the band numbers of recaptured individuals that were first banded on a prior day. We next assessed the age and sex of individuals (Pyle 2022), examined the body and flight feathers for signs of molt, and then determined the WRP code (Pyle 2022; Pyle et al. 2022). We then recorded morphometric information such as fat score, body condition (Kittelberger et al. 2022), wing length, and body mass, the latter of which we calculated by weighing a bird on a scale. Lastly, we noted the net-check time at which an individual bird was captured, along with the specific net in which the bird was found.

### 2.3 Dataset

Since the 2023 solar eclipse occurred on October 14^th^, we filtered our 11 year banding dataset to days that occurred within a two-week period in October, from October 7^th^ to the 21^st^ (hereafter, “study period”). We chose this specific banding window in order to enhance the comparability of banding results that occurred on days around October 14th across multiple years. This choice ensures that data from other banding visits align with a similar timeframe of fall avian movements and includes a similar set of species at this particular location. We also filtered out any captures that did not have the net-check time of capture recorded in our study period. This resulted in 9 birds being removed, of which 3 came from October 7^th^ and 4 came from October 14^th^ in 2020. Additionally, we made two banding visits to the study site in the late afternoon/evening of October 20^th^ and 21^st^ in 2023 (opening at 16:10, closing at 19:10), to record some data on bird movements and responses during low light levels comparable to those experienced during the eclipse.

This resulted in a final Red Butte banding dataset for this study consisting of 22 days of banding in October across 11 years (2012-2023; Table 1), including the 2 evening visits in 2023, and comprised of 292 captures of 22 species (Appendix A). Taxonomic classifications for species followed those of the BirdLife Taxonomic Working Group (BirdLife International 2023), except for Northern Flicker (*Colaptes auratus*) and Yellow-rumped Warbler (*Setophaga coronata*). Subspecies and intergrades were lumped to the species level for Northern Flicker, Dark-eyed Junco (*Junco hyemalis*), White-crowned Sparrow (*Zonotrichia leucophrys*), and Yellow-rumped Warbler. Our dataset for this study includes taxonomic identifications, demographic information (age and sex), capture day and time, and net of capture (Appendix A).

**TABLE 1.** The number of captures of birds across 22 days in our October study period, between 2012 and 2023, while banding in Red Butte Canyon RNA, Utah. The final two visits occurred in the late afternoon to evening.

| **Banding**<br>**Day** | **Number of**<br>**Bird Captures** |
| --- | --- |
| 10/16/2012 | 7 |
| 10/12/2013 | 10 |
| 10/17/2013 | 8 |
| 10/19/2013 | 8 |
| 10/07/2014 | 15 |
| 10/09/2014 | 7 |
| 10/11/2014 | 26 |
| 10/21/2014 | 8 |
| 10/12/2015 | 3 |
| 10/10/2016 | 10 |
| 10/12/2016 | 5 |
| 10/11/2017 | 7 |
| 10/07/2020 | 7 |
| 10/14/2020 | 14 |
| 10/21/2020 | 19 |
| 10/08/2022 | 20 |
| 10/14/2022 | 26 |
| 10/20/2022 | 32 |
| 10/14/2023 | 39 |
| 10/19/2023 | 16 |
| 10/20/2023 | 3 |
| 10/21/2023 | 2 |

### 2.4 Data Visualization

We first plotted the number of bird captures by net-check time for the day of the eclipse. Then, we plotted the trends (mean + 95% confidence intervals) for the day of the eclipse and the overall number of birds captured over time across the 20 days of banding that began in the morning (excluding the two evening banding sessions). For all of these figures, we fit a loess smoothing line (span = 0.6 and 0.7, respectively) to the data to best visualize the trends. All graphing was conducted in R (version 4.3.1, 2023-06-16, R Core Team 2023).

## 3. RESULTS

### 3.1 Solar Eclipse Banding

On October 14^th^, 2023, we arrived at the banding site and opened the nets at 07:30 and closed them at 13:30. The ambient temperature at opening was 38.9°F, and at closing it was 63°F. Throughout the session, a total of 39 individual birds were captured in the mist nets, representing 8 different species. Dark-eyed Junco (*Junco hyemalis*) was the most abundant species, constituting 10 captures (9 Oregon and 1 Slate-colored subspecies), followed by Spotted Towhee (*Pipilo maculatus*; 6 captures), Black-capped Chickadee (*Poecile atricapillus*; 5 captures), and Hermit Thrush (*Catharus guttatus*; 5 captures).

As illustrated in Figure 1, the peak in bird captures occurred at 10:30. The number of species captured exhibited fluctuations corresponding to the changes in sunlight. The morning period maintained a steady capture rate of 2 to 4 individuals per round until 10:00, when 8 individuals were captured, followed by a spike of 11 birds at 10:30 from a single net and then a single capture at 11:00 am, potentially from an earlier entanglement overlooked until after the 10:30 round. Subsequently, there was a return to normalcy from 11:30 to 12:00, with 5 captures, followed by 2 captures at 13:00. The spike of 11 birds at 10:30 consisted of 7 species: 2 Northern Flicker, 2 Ruby-crowned Kinglet (*Regulus calendula*), 1 American Robin (*Turdus migratorius*), 1 Hermit Thrush, 2 Song Sparrow (*Melospiza melodia*), 2 Spotted Towhee, and 1 Oregon Dark-eyed Junco.

**FIGURE 1.**
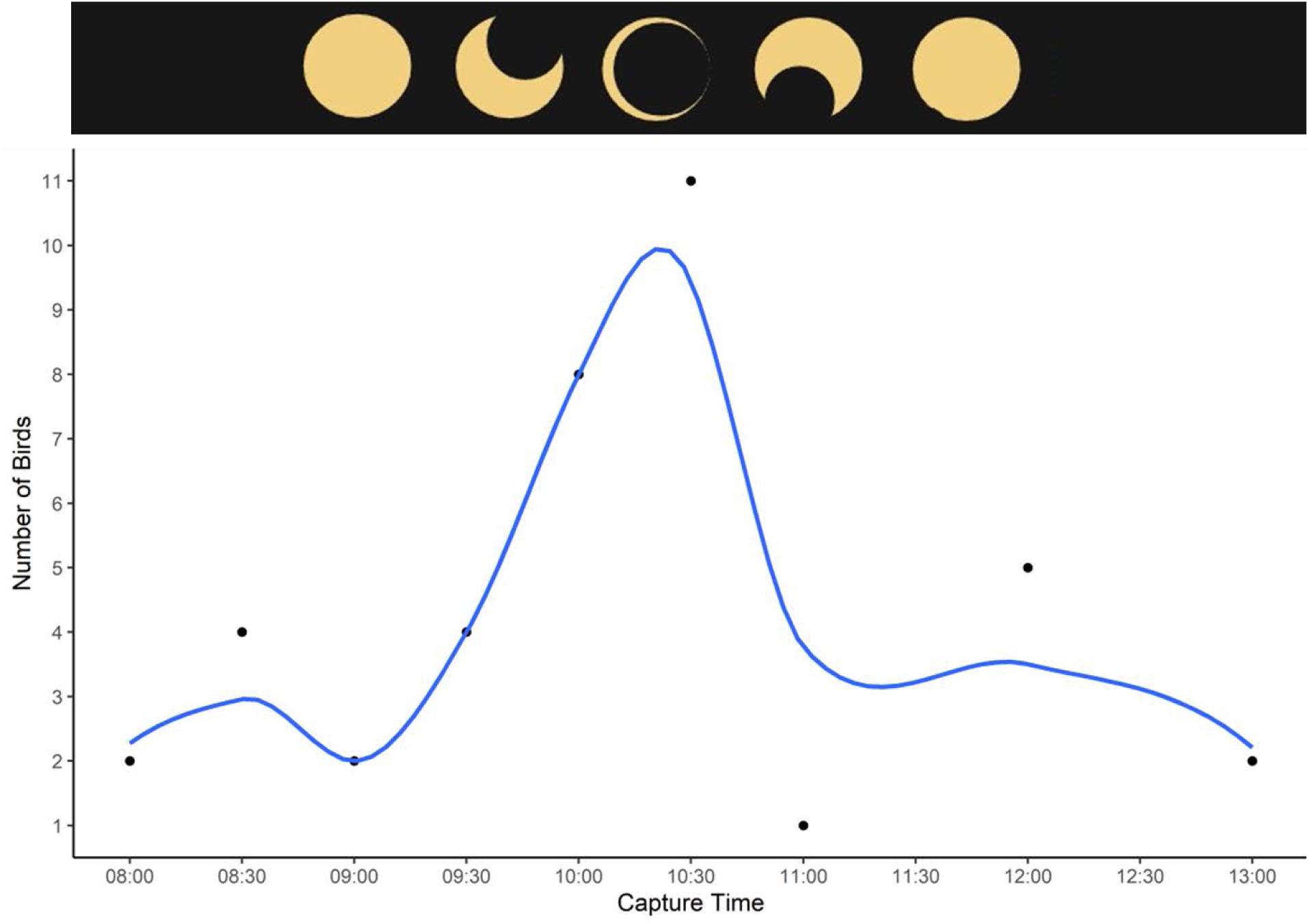
The number of birds captured during different net-check times in Red Butte Canyon RNA, Utah on October 14, 2023, during a partial solar eclipse. A loess smooth line with a span of 0.6 depicts the overall trend in captures. The proportion of totality of the eclipse for different times is noted at the top.

All but two of the birds captured during the main period of the eclipse (check-times 10:00-11:00) were immature, hatch-year birds (the other two individuals were after hatch-year adults), with all the captures on the 10:30 check-time being immatures (Appendix A). However, a majority of the birds captured on this day were immatures, which is to be expected during mid-October (Appendix A). On the other hand, there was a similar mixture of birds of both sexes (along with individuals that were not able to be sexed) captured during the eclipse and throughout the day (Appendix A).

### 3.2 Comparisons with other banding days

The number of birds captured during the eclipse (39) was the most captured on any banding day (April-November) across eleven years at this location (Appendix A). The next closest banding days for number of captures were two visits in mid and late July of 2023, respectively with 34 and 35 captures, at a time when there are many breeding and recently fledged birds in the area. Most of the birds in July are Neotropical migrants that have already migrated from this canyon by mid-October, including many hummingbirds (7 on one of these two days), and only 3 species that we captured on these July dates were also captured during our October study period: Black-capped Chickadee, Song Sparrow, and Spotted Towhee.

During our study period, only one prior banding day, on the 20^th^ in 2022, had more than 30 captures (32); this also was the fourth most numerous banding visit in general at this station. The third and fourth most productive days during this study period were the 11^th^ of 2014 and the 14^th^ of 2022, both with 26 captures (Table 1).

The pattern and peak of birds captured on the day of the eclipse is distinctive from the average trend for captures across all 20 days of morning banding (Figure 2). When looking at the trend from the day of the eclipse with the trends for the other 19 days of morning banding, we can again see that the response of birds during the eclipse is still distinctive from all other banding days in our study period (Figure S1) except for one day: October 11, 2014 (Figure S2). This particular day in 2014 is a bit of an outlier, as the trend is driven by 16 birds captured during the net-check time at 10:00 am, 12 of which came from an apparent mixed-flock of three species flying into the same net (Appendix A). Additionally, the pattern of birds we found during peak totality on the day of the eclipse contrasts sharply with those on our two late afternoon/evening days during the study period, with those days having captures in the low single digits (Table 1).

**FIGURE 2.**
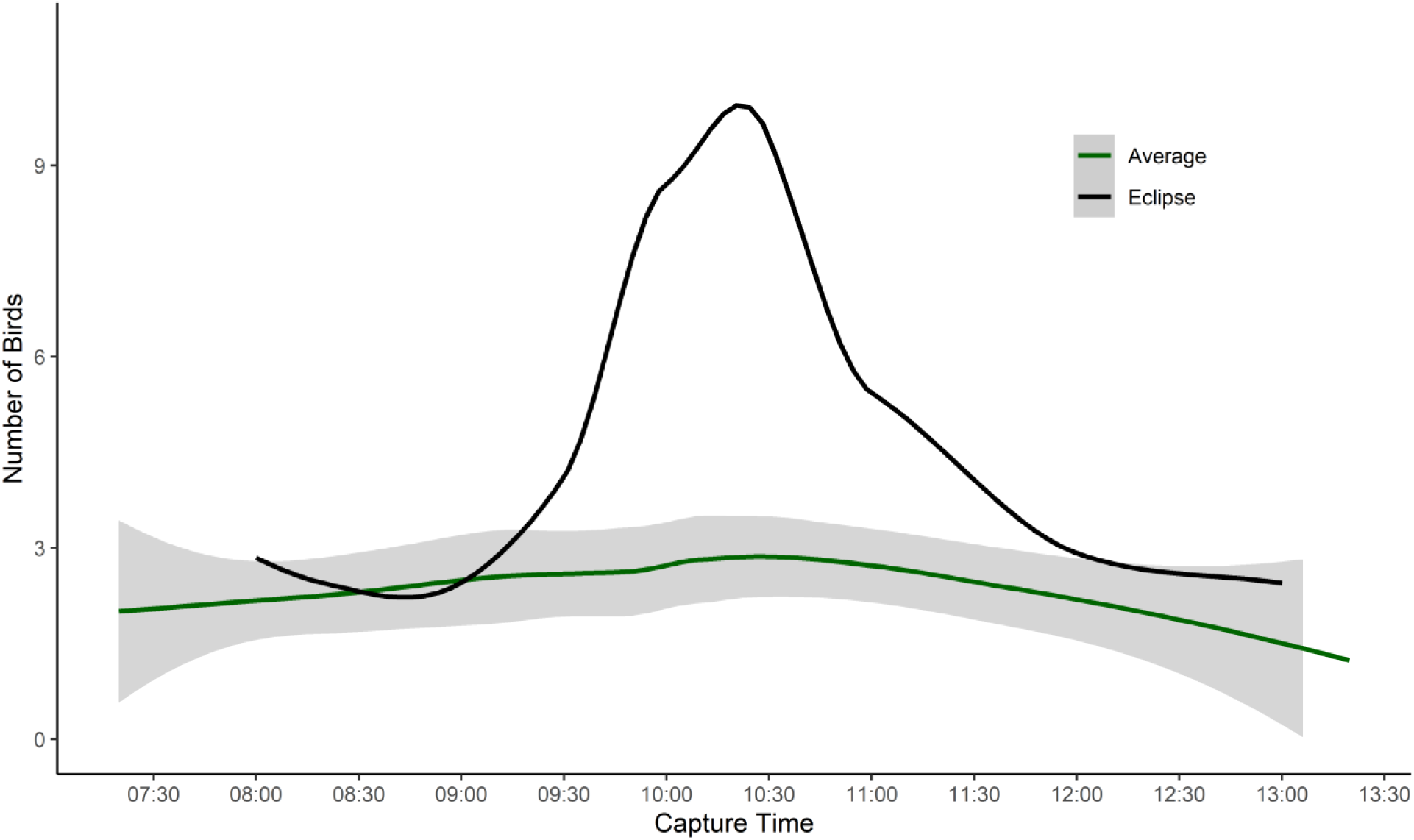
The trend of birds captured during different net-check times on October 14, 2023 (black line), during a partial solar eclipse, compared with the average number of birds captured during 20 total banding visits to Red Butte Canyon RNA, Utah (green line) across an 11 year period (2011-2023). Loess smooth lines with a span of 0.7 are used to best depict the overall trends in captures, and the 95% confidence band is shown for the average trend.

## 4. DISCUSSION

In this study, we used bird banding data to examine changes in bird activity and behavior throughout an annular solar eclipse, comparing the trend from the day of the eclipse with those from prior banding days across an 11-year period. During our banding session on October 14, 2023, we captured 39 individual birds, more than during any other visit to this banding site since 2012. Even though we were not in the path of totality, a majority of the sun was still blocked, with peak obscuration of the sun at our bird banding site reaching approximately 87% (Time and Date 2023). Light levels dropped notably during this period of time, as did the surrounding temperature (pers. observation of the authors). Consequently, we found a strong behavioral response of birds to the onset of the annular solar eclipse, with the eclipse indeed noticeably altering bird activity and movements.

While we originally predicted that bird captures would decline during peak totality, we found the opposite pattern was true. A majority of birds (24 individuals, 62% of those banded this day) were captured during the main period of the eclipse (between 9:30 and 11:30 am), with the biggest spike in bird captures coinciding with the peak of partial totality around 10:30 am (Figure 1). Even though we captured one bird on the following round at 11:00 am (Figure 1), it is very likely that this Spotted Towhee had become entangled during a prior round and just was not reached until later, since the net preceding the towhee had 11 birds in it. After the eclipse finished, bird captures returned to a normal baseline level (Figure 1). Furthermore, while we visited the banding site on a couple evenings after the eclipse to assess movements of birds in low light levels comparable to those on October 14^th^, we found that very few birds were moving during those evening sessions, supporting that the spike in bird captures during the eclipse was a result of birds responding to the anomalous nature of the sun being obscured in contrast to the sun regularly setting for the day. The trend of bird captures during the eclipse also notably differs from the average trend during mid-October morning banding sessions across the past 11 years (Figure 2, S1), as well as the trend for 18 other banding days during our study period (Figure S2), suggesting the bird movements we recorded on October 14^th^ are atypical for this location at this time of year and again an artifact of the eclipse.

The positive relationship we found in bird movements during the eclipse runs counter to behavioral responses of birds in prior studies, which noted declines in overall activity and movements of birds during eclipses (Tramer 2000; Nilsson et al. 2018; Wiantoro 2019; Mekonen 2021). Three of these studies though were observational in nature rather than relying on data collected from a rigorous methodology (Nilsson et al. 2018), like bird banding. Birds may have been imitating their typical end-of-day behaviors when confronted with the sudden onset of darkness from an eclipse. As peak totality approached, birds could have been moving around more in order to return to or find a place to roost (Van Doren et al. 2017; Nilsson et al. 2018; Mekonen 2021), thinking that dusk was occurring; the mist-nets therefore could have prevented these birds from reaching intended roost spots to remain relatively inactive for the remainder of the eclipse, in which case trends in activity would have been more similar to those found in other studies. On the other hand, the unusual nature of the eclipse occurring during the day could have caused birds to become disoriented and move around in a chaotic and stressed manner. This may be supported by the fact that there was a community-level response of birds to the eclipse, with a mixed-species assortment of birds captured during the height of totality that consisted of 7 species in a single net (Appendix A). Regardless, the trend in movements we found suggests a consistent disruption in avian circadian rhythm (Hartstone-Rose et al. 2020), where alterations in light (and possibly temperature) triggered a change in movement.

Additionally, while we saw an increase in bird captures as the eclipse proceeded, we observed a negative relationship with the avian soundscape during the eclipse: it became very quiet, with seemingly no birds vocalizing besides a Northern Flicker, which was actually calling from the net in which it had flown. Birds had been vocalizing before the light levels dropped, and once the sun became unobscured again, the soundscape returned to normal (pers. observation of the authors). This decrease in avian vocal activity during the height of the eclipse mirrors trends reported in some prior studies (Mendoza 2018; Mekonen 2021), but differs from another that saw an increase in vocalizations (Brinley Buckley et al. 2018). These differences in vocal responses could be driven by phylogeny and species-specific behavioral cues for vocalizing (Brinley Buckley et al. 2018), with some species more inclined to vocalize than others in response to photic disturbances or lower light levels. However, one of the species noted in Brinley Buckley et al. (2018) that increased its vocalizations during an eclipse, Song Sparrow, was also captured during our eclipse banding session and was not discernibly vocalizing during totality. The eclipse in 2017 occurred in mid-August though, during the end of or after the breeding season when birds are still fairly vocal, and the species that showed significant increases in vocalization were late-season breeders (Brinley Buckley et al. 2018); on the other hand, the eclipse in 2023 occurred in mid-October, during fall migration when birds are less vocal. The seasonal timing of when an eclipse occurs may therefore have an important effect on the vocal responses of some species.

Besides seasonal timing, our findings may not be broadly applicable to those from different geographical locations, different bird species, or other solar eclipses. The specific conditions of this eclipse (e.g., path of totality, percentage of obscuration) may influence bird behavior differently than other eclipses. In addition, this study is based on observations during a single annular solar eclipse on October 14, 2023, in Salt Lake City, Utah. Therefore, our study focuses on a relatively small dataset, consisting of observations from one specific location and one eclipse event. The limited sample size may affect the robustness and generalizability of the findings. Continued work on this topic would benefit from including data from multiple eclipse events and locations to strengthen the conclusions. To accomplish this in the future, researchers in North America could take advantage of the upcoming total solar eclipse on April 8^th^, 2024. Replicating this study in banding stations close to or within the path of totality on this occasion could enhance the generalizability of the results, shedding light on potential patterns or variations in avian responses across multiple eclipse events and reinforcing the robustness of the study’s methodology in different eclipse scenarios.

## 5. CONCLUSION

This study marks the first known published effort to conduct bird banding during a solar eclipse, offering important insights into the behavioral responses of birds to changes in light conditions during this infrequent celestial occurrence. We documented a notable increase in bird movements before and during the height of the eclipse, which we may not have been able to detect if we had not been using nets to capture birds as they moved through the vegetation and habitat and instead been relying on visual observations or changes in the avian soundscape. By leveraging bird banding data from the 2023 solar eclipse, compared with data from a similar period of time from the same site over the past 11 years, we provide a unique perspective on avian behavior during celestial events, moving beyond anecdotal evidence and contributing scientific rigor to the field. Emphasizing the significance of bird banding as a methodological approach, the study conducts a quantitative analysis of bird captures, systematically examining species composition and capture trends, thus providing a robust and data-driven understanding of avian responses to the eclipse. This study paves the way for a more comprehensive understanding of how birds respond to celestial events, and also provides a foundation that future researchers and bird banders can build off of during other solar eclipses.

## 6. CONFLICT OF INTEREST

The authors declare that the research was conducted in the absence of any commercial or financial relationships that could be construed as a conflict of interest.

## 7. AUTHOR CONTRIBUTIONS

KDK and AD wrote the manuscript; AD, AMS, and KDK conceived the original idea for the project; KDK managed the banding site, with support from ÇHŞ; AD and KDK prepared the banding dataset; KDK plotted the data and created the figures, with assistance from AD; all co-authors contributed to and gave final approval of the manuscript.

## 8. ACKNOWLEDGEMENTS

We thank Flávio Mota and Nicholas Seefeldt for joining us banding on the evenings in October 2023, and prior volunteers that banded on other days used in this study. We also thank Asja Johnson and Amy Buxton for entering and proofing the 2023 data, and to the countless banders and volunteers over the years that have helped collect this data. We also thank Colby Tanner for providing feedback on visualizing the data and thoughts on the manuscript itself.

*FUNDING* We are grateful to H. Batubay Özkan and Barbara Watkins for their support of the Biodiversity and Conservation Ecology Lab at the University of Utah, School of Biological Sciences. We are grateful to the University of Utah’s Graduate Research Fellowship for providing support to the lead author to carry out this research. We are grateful to the Medium Grant from the Sustainable Campus Initiative Fund (SCIF) at the University of Utah for helping fund equipment used during banding seasons.

## 9. DATA AVAILABILITY

The dataset of bird captures used in this study are available in Appendix A.

## Notes

### Competing Interest Statement

The authors have declared no competing interest.

