## Supplemental Figures 1 and 2 for "Solar Bird Banding: Notes on Changes in Avian Behavior While Mist-netting During an Eclipse"

**FIGURE S1**. Trends of birds captured during different net-check times across 19 banding days (October 11, 2014 is removed) in mid-October in Red Butte Canyon RNA, Utah across an 11 year period (2011-2023). The trend for birds during the partial solar eclipse on October 14, 2023 is shown in a black line, with the average number of birds captured during 19 total banding visits depicted in green. Loess smooth lines with a span of 0.7 are used to best depict the overall trends in captures, and the 95% confidence band is shown for the average trend.

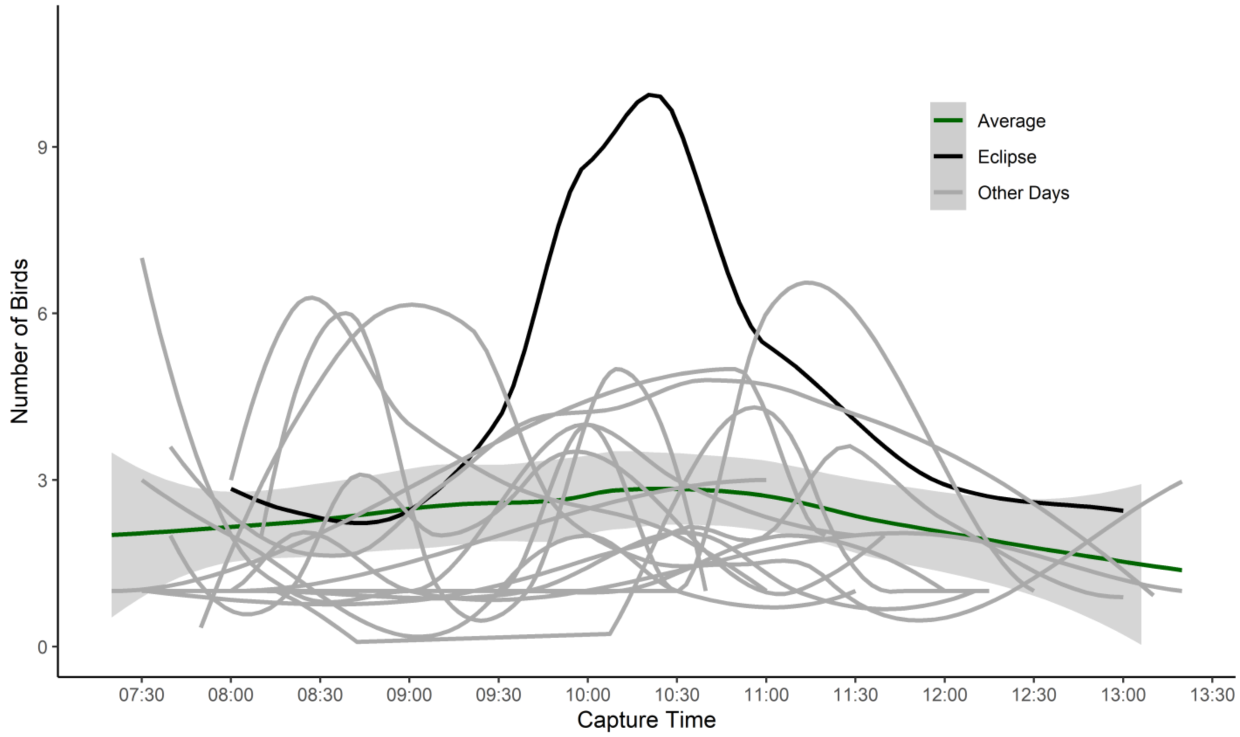

**FIGURE S2**. Trends of birds captured during different net-check times across 20 banding days in mid-October in Red Butte Canyon RNA, Utah across an 11 year period (2011-2023). The trend for birds during the partial solar eclipse on October 14, 2023 is shown in a black line, the trend for outlier day October 11, 2014 is in blue, and the average number of birds captured during all the banding visits is depicted in green. Loess smooth lines with a span of 0.7 are used to best depict the overall trends in captures, and the 95% confidence band is shown for the average trend.

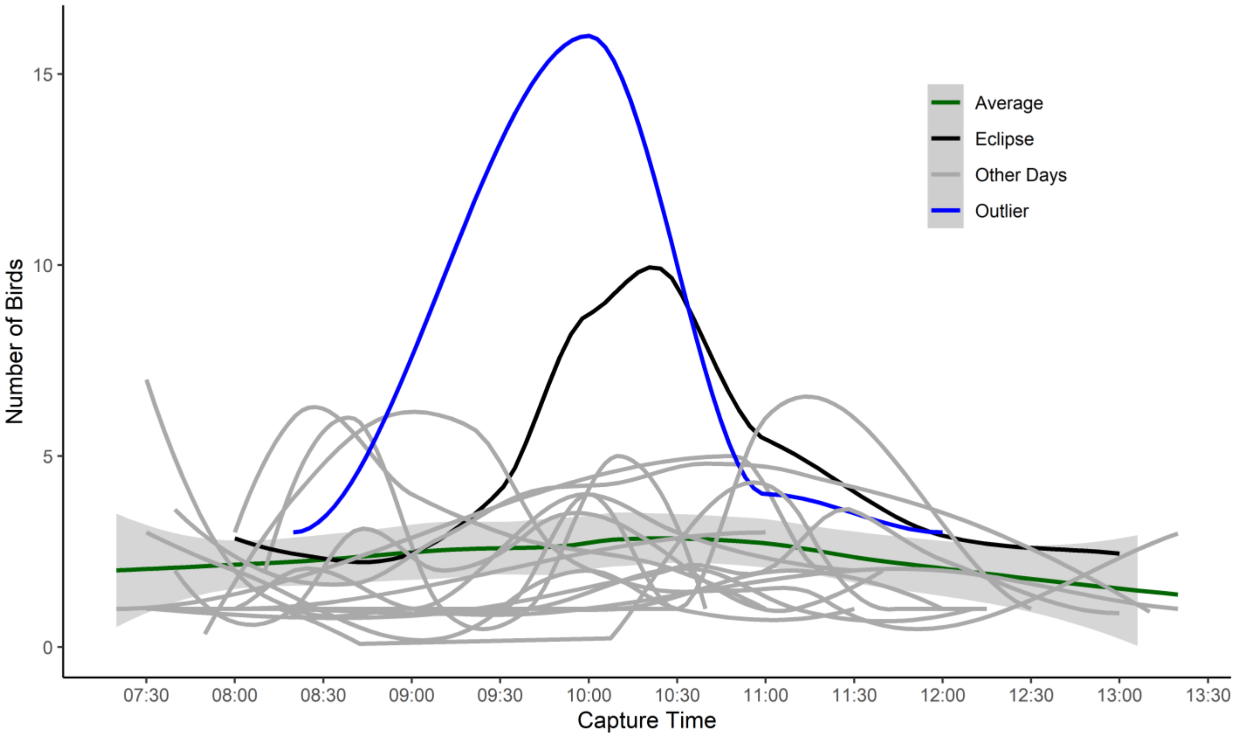
